# Meiotic Drive and Survival Probability of Newly Inverted Chromosomes

**DOI:** 10.1101/486712

**Authors:** Spencer A. Koury

## Abstract

When a new gene arrangement is generated by spontaneous mutation its survival is uncertain and largely unaffected by associated fitness effects. However, if a new chromosomal inversion is introduced into a population already polymorphic for inversions, then its survival probability will be a function of the relative size, position, and linkage phase of the gene rearrangements. This dependence on structural features is due to the complex meiotic behavior of overlapping inversions generating asymmetric dyads, which in turn cause both underdominance and meiotic drive/drag. Therefore, survival probabilities of new inversions can be expressed in terms of the probability of forming an asymmetric dyad via crossing over in meiosis I and the probability of recovery from that asymmetric dyad during disjunction in meiosis II. This model of female meiotic drive was parameterized with data from published experiments on laboratory constructs in *Drosophila melanogaster*. Generalizing this analysis to all possible inversions predicts a bias towards larger, proximally located inversions having a shorter persistence time in populations. These population genetic predictions are consistent with cytological evidence from natural populations of *D. melanogaster*. This research mathematically formalizes a cytogenetic mechanism for female meiotic drive/drag as the major force governing behavior of new gene arrangements entering populations, and therefore determining the genomic distribution of segregating inversion polymorphism.

## Introduction

The cytological discovery of polytene chromosomes in salivary glands of *Drosophila* larvae greatly expanded the study of classical genetics (Painter 1933, Sturtevant 1965). For cytogenetics, the banding pattern of polytene chromosomes provided a physical basis for genetic maps of *Drosophila melanogaster* and direct observation for a vast array of chromosomal aberrations (Bridges 1935, Sturtevant and Beadle 1939). For evolutionary genetics, the banding pattern confirmed conservation of euchromatic chromosome arms known as Muller elements and allowed easy genotyping of chromosomal inversions in virtually any *Drosophila* species (Muller 1940, Sturtevant and Novitski 1941). What followed was an explosion of research documenting inversion polymorphism in populations of hundreds of different species, the first phylogeny based on genetic data, and the start of the widely acclaimed *Genetics of Natural Population Series I-XLIII* (Sturtevant and Dobzhansky 1936, Dobzhansky and Queal 1938, Provine 1981, Sperlich and Pfriem 1986).

The extensive documentation of inversions in natural populations of *Drosophila* species, especially in the *obscura* group, has allowed fundamental insights into the origins, establishment, and ultimate loss or fixation of inversions (Dobzhansky and Epling 1944, Wallace 1953, Ohta and Kojima 1968, Lande 1984, Schaeffer 2008). The distribution of inversion breakpoints along a chromosome has been particularly informative, as these breakpoints exhibit several nonrandom patterns that have inspired models of inversion evolution (Federer et al. 1967, Van Valen and Levins 1968, Krimbas and Powell 1992). One class of models explains the distribution of inversion breakpoints in terms of selection for inversion length or chromosomal placement and the resulting recombination suppression (Caceres et al. 1997, Cáceres et al. 1999, Corbett-Detig 2016). Alternately, inversion breakpoint patterns have been attributed to non-selective mechanisms by: 1) modeling the spontaneous mutational process as transposable element mediated, 2) using directionally biased spontaneous mutation rates, or 3) considering structural heterozygosity as autocatalytic, such that pairing difficulty at breakpoints causes chromosomal breakage and elevates rates of gene rearrangement (Novitski 1946, Bernstein and Goldschmidt 1961, Lim and Simmons 1994). Finally, a meiotic drive mechanism has been proposed to explain the distal shift in breakpoints for serially inverted chromosomes. There is no consensus on what balance of evolutionary forces govern the fate of inversion polymorphism; however, the vast majority of research on inversion evolution is either directly or indirectly based on the fundamental observation that chromosomal inversions in the heterozygous state suppress recombination (Krimbas and Powell 1992, Hoffmann and Rieseberg 2008).

In absence of recombination the constant input of deleterious mutations causes the successive loss of the most fit chromosomes and delays the removal of deleterious mutations from populations (Muller 1964). Ohta and Kojima (1968) modeled this process for inversions with a time heterogeneous branching process, and concluded the ultimate extinction of an inversion is certain unless unique mechanisms intervene to maintain selective superiority. Selection on epistatic fitness effects unique to the inversion may be one such mechanism (Dobzhansky 1948, Ohta and Kojima 1968, Feldman et al. 1996). If such epistatic fitness effects exist (e.g. Schaeffer et al. 2003, Houle and Márquez 2015), the rate at which the multi-locus combination of favorable alleles is broken up would be reduced in inverted regions. Thus, inversions create conditions favorable for both the irreversible mutational decay of fitness, as well as the build-up and preservation of co-adapted gene complexes. This double-edged sword is responsible for the observation that inverted chromosomes sampled from natural populations carry 1.5 times as many lethal mutations and have only 78% the homozygous viability of standard arrangement chromosomes (Mukai and Yamaguchi 1974), while at the same time being associated with local adaptation (Schaeffer 2008, Lowry and Willis 2010, Jones et al. 2012), coadapted gene complexes (Jaenike 2001, Lyon 2003, Larracuente and Presgraves 2012), evolution of neo-sex chromosomes (Charlesworth et al. 2005, Bachtrog 2013, Tuttle et al. 2016), and speciation (White 1978, Noor et al. 2001, Rieseberg 2001).

Recombination suppression for inversions heterozygous with the standard arrangement is the combined effect of reduced crossing over in inverted regions and elimination of acentric and dicentric products of single crossover events (Sturtevant and Beadle 1936, Novitski and Braver 1954). In higher Dipterans the symmetrical meiosis in males is achiasmate (for rare exceptions see Gethmann 1988), meaning that inversion heterozygosity in males can have no effect on recombination. Female meiosis is asymmetric, for every oocyte only one of the four meiotic products results in a functional gamete. In *Drosophila*, female meiosis arrests in metaphase of meiosis I and only goes to completion after fertilization (King 1970). Prior to metaphase I the four meiotic products are arranged in a linear array such that only the terminal position migrates to the egg pole (Huettner 1924). Because single crossovers in inverted regions cause dicentric chromosomes that are mechanically constrained to occupy the two internal positions, they are never included in the functional egg (Sturtevant and Beadle 1936). The structures caused by homologous pairing and resulting crossover products of female meiosis are illustrated in figure 1a in absence of inversions, figure 1b in heterozygotes with a single inversion, and figure 1c in heterozygotes with two overlapping inversions.

**Figure 1.**
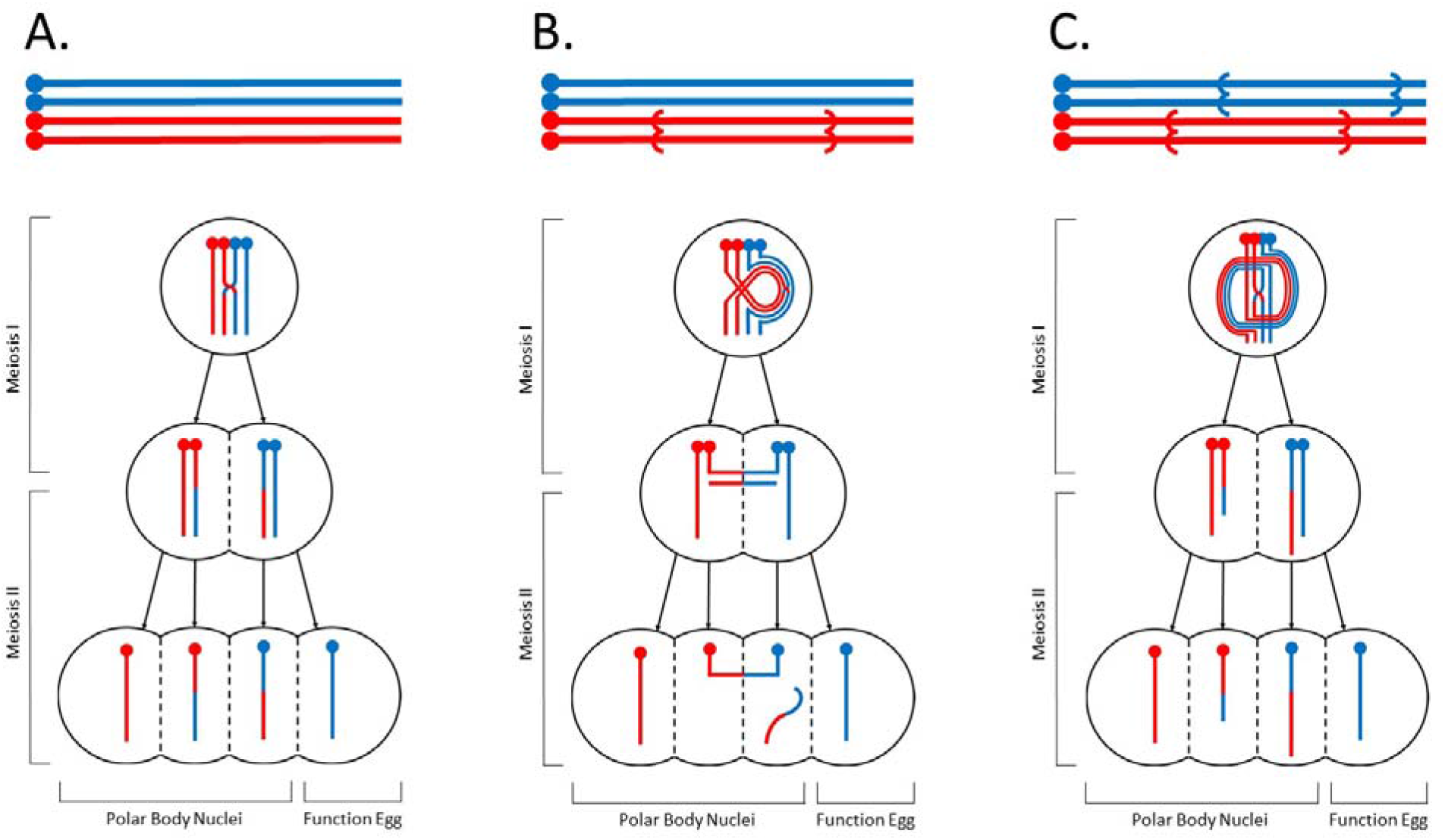
Pairing diagram for inversions, synapsis, and the resulting crossover products in the context of a) homokaryotypes with no inversions, b) heterokaryotypes with a single inversion, and c) heterokaryotypes of two overlapping inversions. Note all crossover products between overlapping inversions are monocentric but not of equivalent length, forming asymmetric dyads.

The acentric-dicentric elimination mechanism described above cannot apply to crossing over in the homosequential regions (*i.e*. shared inverted regions) of heterozygotes for two overlapping inversions. Because gene order is the same in the shared inverted regions of overlapping inversions, crossing over creates four monocentric meiotic products (figure 1c). Although no dicentric chromosomes are formed, the inverted regions that are *not* shared between overlapping inversions are either duplicated or deleted. The aneuploidy generated by crossing over has two consequences: 1) large duplications and deletions are dominant lethals, and 2) size asymmetry of the chromatids in each dyad results in unequal transmission. The latter effect is known as non-random disjunction, a form of female meiotic drive that has been extensively studied using compound chromosomes in *D. melanogaster* (Novitski and Sandler 1956, Sandler 1958, Sandler and Lindsley 1963, Lindsley and Sandler 1965, Lucchesi 1965, Sandler 1965, Sandler and Lindsley 1967).

This meiotic drive mechanism was recently invoked to explain a distal shift of inversion breakpoints in the evolution of serially inverted chromosomes of *D. obscura* group species. As previously outlined, single crossovers in the shared inverted regions (or in serially inverted regions) create asymmetric dyads. Experiments with compound chromosomes, translocation heterozygotes, overlapping inversions and heterochromatin displacement all confirmed that the smaller chromatid of an asymmetric dyad is preferentially included in the functional egg (Novitski 1951, Novitski 1967). As a consequence of 1) the pairing geometry of overlapping inversions, 2) the formation of asymmetric dyads, and 3) the preferential recovery of shorter chromatids, the distal inversion in heterozygotes for overlapping inversions will always *drive* and the proximal inversions will always *drag*. Although this mechanism was initially studied in relation to an obscure pattern of serially inverted chromosomes, it can also shed light on several better known and similarly unexplained patterns of inversion polymorphism. For example, the subtelomeric concentration of inversions, Wallace’s rule of triads, the autocorrelation of inversion breakpoints, and the intraspecific length of gene arrangement phylogenetic series are all qualitatively consistent with this meiotic drive mechanism favoring distal inversions.

The purpose of the present study is to parameterize the model of nonrandom disjunction for overlapping inversions and quantify the population consequences of this form of meiotic drive/drag. First, the branching process was used to calculate the survival probability of a spontaneous gene rearrangement in a monomorphic infinite population of diploid individuals under mendelian segregation. Second, expressions for the transmission bias and frequency of lethal zygotes produced by females heterozygous for overlapping inversions were given in terms of the probability of forming an asymmetric dyad and the probability of recovery from an asymmetric dyad. Third, the survival probability for spontaneous inversion mutations in a polymorphic population was modeled by incorporating transmission bias and underdominance (frequency of lethal zygotes). Parameter estimates for the meiotic events and the resulting fitness effects were derived from published cytogenetic experiments in *D. melanogaster*. This parameterization allowed calculation of survival probabilities for all possible inversions entering a natural population of *D. melanogaster*. Converting these survival probabilities to persistence (time to extinction) and pervasiveness (total number of individuals affected) predicts biases in the genomic distribution of segregating inversion polymorphism. Finally, the biases in inversion size, position, and phase generated by this cytogenetic mechanism were compared to empirical data on inversion polymorphism in natural populations of *D. melanogaster*.

### Survival Probability in a Monomorphic Population

The probability that a mutant gene survives to the next generation is a classical problem in population genetics and was first addressed by Fisher (1923). The branching process treatment formalized by Haldane (1927), considers the fate of a mutant gene represented by a single copy in a population of infinite size. Rather than follow changes in allele frequency this model generates probability distributions for the absolute number of copies of the mutation (Crow and Kimura 1970). In contrast to the textbook example, this study examines the branching process in diploid sexually reproducing organisms, making the probability of surviving to the next generation subject to both the demographic effect of stochastic variation in family size and the stochastic processes inherent to transmission genetics of diploids. Assuming family size (*n*) is Poisson-distributed with mean size (*c*) and the mutant gene’s transmission probability (*k*) is described by the binomial distribution, then the probability of having *m* copies of the mutant gene in the first generation is:

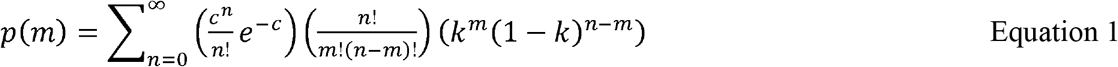

The branching-process uses this single generation distribution of gene copies to describe the forward time behavior of the mutant with a recursion equation. Determining the fate for a newly mutated chromosomal rearrangement, and the timing of that fate, is reduced to calculating the probability of zero copies in generation *t*. The probability-generating function for this distribution is *g*(*x*) = *e*^*ck*(*x*−1)^. After appropriate substitution (*cf*. Crow and Kimura 1970 pg. 421) the probability of surviving (*u*) to generation *t* is:

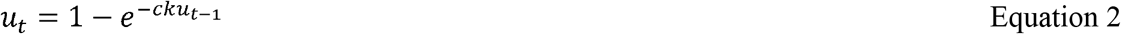

This equation can be applied to the simple case of a spontaneous gene rearrangement in a monomorphic population at equilibrium and in absence of any fitness effect (*c = 2*). If this mutant allele segregates according to Mendel’s second law (*k = 0.5*), then this equation simplifies to the classic model of Haldane (1927). However, the expanded form of the recursion equation is necessary for more complex situations where variation in average family size (*c*) and moreover transmission rate (*k*) will be directly affected by cytogenetic mechanisms. The meiotic behavior of overlapping inversions described below presents just such a situation.

#### Nonrandom Disjunction of Overlapping Inversions

In heterozygotes for overlapping inversions there always exists a shared inverted regions whose gene order is homosequential. Crossing over in this homosequential region is not suppressed by the usual mechanism preventing recombination amongst chromosomal rearrangements (*i.e*. failure of proper segregation of acentric and dicentric chromosomes). In heterozygotes for overlapping inversions, all four possible products of female meiosis will be monocentric and therefore could in principle be included in the functional egg. However, classical experiments from *Drosophila* cytogenetics predict these four meiotic products will not have the same probability of migrating to the egg pole (Glass 1934, Sturtevant and Beadle 1936, Novitski and Braver 1954, Zimmering 1955, Lindsley and Sandler 1965). The unequal recovery of complementary meiotic products in these situations is a well-established phenomenon termed nonrandom disjunction and has been thoroughly reviewed (Novitski 1951, Novitski 1967).

The strength of transmission bias due to nonrandom disjunction for overlapping inversions can be modeled in terms of fundamental processes of crossing over in meiosis I and disjunction in meiosis II. The four possible meiotic products are chromosomes with the distal inversion, duplications, deletions, and the proximal inversion (Figure 1c). The geometry enforced by homologous pairing of overlapping inversions requires that the chromatid carrying crossover-generated large duplications is always paired with the distal inversion in an asymmetric dyad entering meiosis II. Conversely, the recombinant chromatid carrying the large deletions is always paired with the proximal inversion in meiosis II. There are no possible exceptions to this rule of pairing the larger crossover product with distal inversions and the smaller crossover product with proximal inversions. From each of these asymmetric dyads the shorter chromatid has a greater chance of being included in the functional egg (Novitski 1951, Novitski 1967). As a consequence, in females heterozygous for overlapping inversions the distal inversion will always drive while the proximal inversion will always drag.

If the probability of creating an asymmetric dyad from crossing over in the shared inverted region is *a*, and the probability of recovering the shorter chromatid from an asymmetry dyad is *r_i_*, with *i* specifying the dyad containing either the proximal or distal inversion, then the frequency with which the meiotic products are included in the functional egg are given in table 1a. Because large duplications and deletions cause dominant lethal effects, females heterozygous for overlapping inversion are expected to exhibit fitness underdominance. If the aneuploid classes causing lethal zygotes are combined, as are non-recombinant and recombinant classes for the chromatids carrying proximal and distal inversions, then the frequency of meiotic products in the functional eggs of a female heterozygous for overlapping inversions is given in table 1b. From these frequencies the underdominant effect on family size is expressed as:

**Table 1.** a) The frequency of all possible meiotic products as a function of the probability of crossing over (a) and recovery (r) in the functional egg. b) The frequency transmitting a meiotic product carrying either a distal inversion, lethal mutation, or proximal inversion as a function of the probability of crossing over (a) and recovery (r) in the functional egg.

| a) |  | b) |  |
| --- | --- | --- | --- |
| Class of Meiotic Products | Frequency in Functional Egg | Class of Meiotic Products | Frequency in Functional Egg |
| Non-recombinant Distal Inversion | $(1/2) (1-a)$ | Distal Inversion | $\frac{1 - a + ar_{dist}}{2}$ |
| Recombinant Distal Inversion | $(1/2) (a) (r_{dist})$ | Lethal Mutation | $\frac{a(1-r_{dist}+r_{prox})}{2}$ |
| Recombinant Double Duplication | $(1/2) (a) (1-r_{dist})$ | Proximal Inversion | $\frac{1 - ar_{prox}}{2}$ |
| Recombinant Double Deletion | $(1/2) (a) (r_{prox})$ | | |
| Recombinant Proximal Inversion | $(1/2) (a) (1-r_{prox})$ | | |
| Non-recombinant Proximal Inversion | $(1/2) (1-a)$ | | |

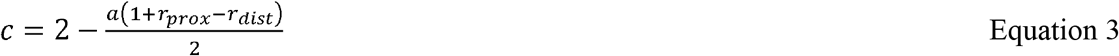

The transmission bias amongst viable progeny favoring the distal inversion *(i.e*. meiotic drive) is:

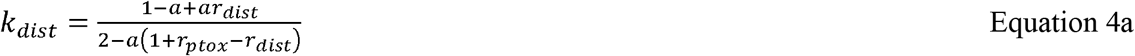

Conversely, the undertransmission (*i.e*. meiotic drag) of the proximal inversion amongst viable progeny can be expressed as:

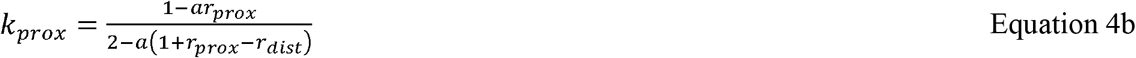

This section’s treatment and illustrations implicitly considered overlapping inversions on opposing chromosomes (repulsion phase). However, the exact same mechanism operates for overlapping inversions on the same chromosome (coupling phase), such as those observed in serially inverted chromosomes (supplemental figure 1a). Similarly, if the new inversion is entirely included in the inverted region, rather than overlapping the segregating gene rearrangement, then meiotic drive/drag will occur in both coupling and repulsion phase (supplemental figure 1b and 1c). There are, however, two salient modifications for these scenarios. First, inversions in coupling phase drive/drag only when heterozygous with the standard arrangement. Second, the magnitude of drive/drag for included inversions is reduced because both aneuploid meiotic products carry one duplication and one deletion rather than one meiotic product carrying two duplicated regions or two deleted regions as is the case for overlapping inversions. As noted previously, there are no possible exceptions to the pairing rule forming asymmetric dyads of larger crossover products with the distal inversion, even if the larger crossover product contains both a duplication and a deletion.

In summary, whenever two different but overlapping gene rearrangements of the same chromosome arm are segregating in a population, meiotic drive/drag is predicted with the strength of transmission bias depending on relative size, position, and linkage phase of the inversions. The strength of drive can be modeled with two parameters, the probability of forming an asymmetric dyad (*a*) and the probability of recovery from that asymmetric dyad (*r_i_*). In following sections, the effects of this cytogenetic mechanism will be incorporated in the survival probability recursion equation, where the importance of relative size, position, and linkage phase of inversions in modifying the cytogenetic mechanism will be considered. Finally, parameter estimates will be provided for nonrandom disjunction in *D. melanogaster* that allow the population consequences of the meiotic drive/drag mechanism to be quantified.

### Survival Probability in a Polymorphic Population

When a spontaneous gene rearrangement occurs in a population already polymorphic for chromosomal inversions its survival probability is altered by the complex meiotic behavior of overlapping inversions. As demonstrated in the previous section, the fitness effects and transmission bias generated during meiosis can be expressed as functions of the probability of crossing over and recovery from an asymmetric dyad (equations 3 and 4a,b). Therefore, the survival probability recursion equation is modified by incorporating equation 3 and equation 4a or 4b depending the relative position of the new inversion, yielding:

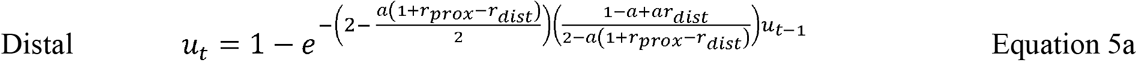

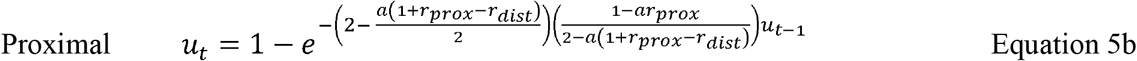

As previously noted, these effects are both sex-specific and genotype-specific; first, recombination and asymmetric meiosis is limited to females in *Drosophila*, and second, drive/drag occurs only for inversions in repulsion phase or for serially inverted chromosomes when heterozygous with the standard arrangement. As a consequence, further description of the modeled population is necessary.

A discrete generation model was used to calculate survival probabilities of a newly mutated chromosomal inversion during the early phases of establishment. This is a model of a dioecious population of infinite size, but with countable individuals. This is a random mating population with equal sex ratios. In this population there is a common chromosomal inversion segregating for one of the autosomes at frequency *q*. If we introduce a second inversion by mutation to this same autosome, then the survival probabilities of this new chromosomal rearrangement can be modeled by the branching process.

The infinite, but countable, assumption has two important consequences for a new mutation. First, the new chromosomal rearrangement will always be found in the heterozygous state independent of the number of copies that exist in the population. Second, allele frequencies of other chromosomal arrangements are unaffected by changes in the count of the new chromosomal rearrangement. Therefore, a spontaneous rearrangement occurs in a context capable of meiotic drive/drag with frequency 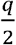 if it is repulsion phase with the older segregating inversion, and 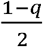 if it is in coupling phase with the older inversion (i.e. a serially inverted chromosome). The remainder of the time the new inversion is either in a heterozygous state with an arrangement against which it cannot drive/drag or in a male where recombination does not occur. Applying the appropriate weighting to each scenario for a new inversion distally positioned in repulsion phase (equation 6a), proximal repulsion phase inversions (equation 6b), distal coupling phase inversions (equation 6c), and proximal coupling phase inversions (equation 6d), then the survival probabilities become:

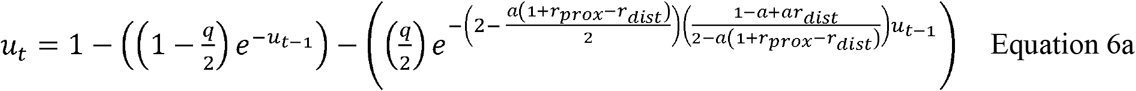

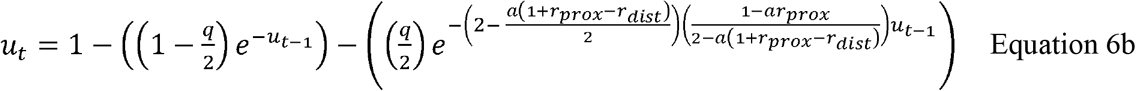

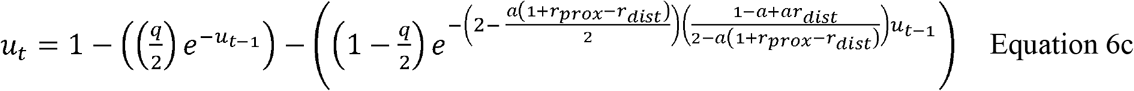

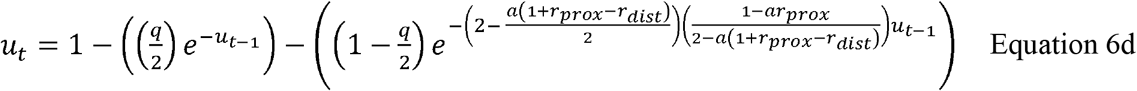

### Parameterization of Nonrandom Disjunction

The meiotic drive/drag model presented is general for any organism with the following three conditions: 1) asymmetric female meiosis, 2) crossing over in heterokaryotypes during meiosis I, and 3) non-random disjunction from asymmetric dyads in meiosis II. While these conditions are likely to be very common, experimental analysis of the second and especially the third conditions are largely limited to *D. melanogaster*. The estimates presented here are a first attempt at parameterizing this model for inversion evolution based on published experimental data. This parameterization comes with the caveat that certain aspects may be species-specific and subsequent revision may be required when adapting this framework to the vagaries of meiosis in new model systems.

Crossing over occurs in heterozygous inverted regions, however, estimation of these rates is difficult and relies on either indirect measures or complex experimental constructs. Rates of single crossover can be inferred from reduced viability of progeny from females heterozygous for pericentric inversions (Coyne et al. 1991, Coyne et al. 1993, Navarro and Ruiz 1997). Alternately, compound chromosomes can be used to recover the crossover products from females heterozygous for paracentric inversions usually eliminated as acentric or dicentric chromosomes (Sturtevant and Beadle 1936, Novitski and Braver 1954, Hinton and Lucchesi 1960). Finally, in special cases Sturtevant and Beadle (1936) were able to collect recombinant chromosomes from overlapping inversions by outcrossing to translocation stocks that complemented crossover-generated deficiencies. All three methods require extensive experimental control and statistical correction for both viability effects and presence of meiotic drive (Novitski 1951). Consensus results reveal crossing over is reduced in heterozygous inverted regions, but this reduction is dependent on the corresponding genetic length of the inverted region on the standard genetic map (figure 2a,b,c). The probability of crossing over (*a*) is modeled as a function of the size of the inverted region (*s*) corrected by the fraction of the total genetic length of the chromosome (*l*) that is inverted.

**Figure 2.**
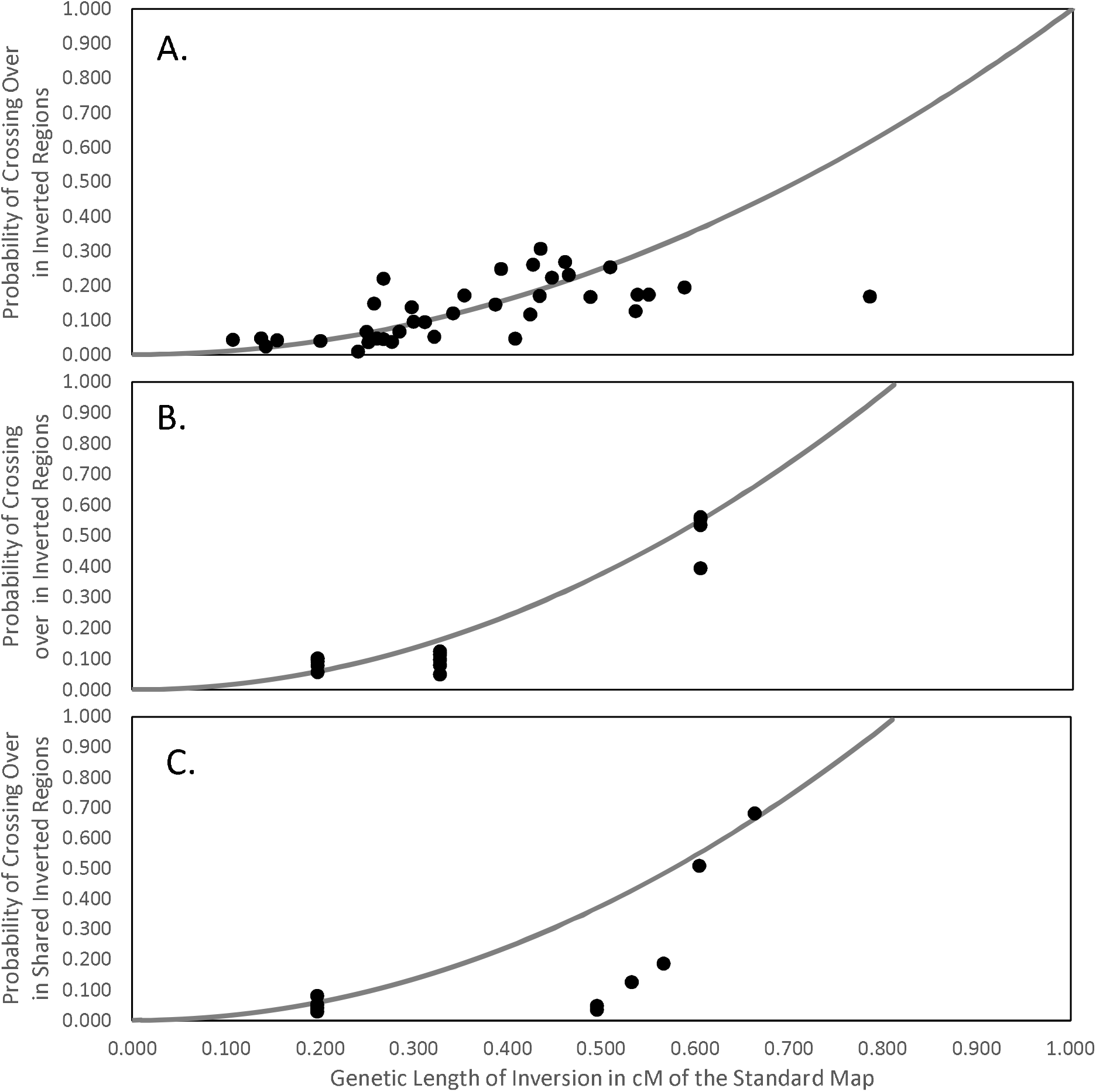
The probability of crossing over in a) inverted regions of pericentric inversions, b) the inverted regions of paracentric inversions, and c) the shared inverted region of overlapping paracentric inversions. The illustrated function is equation 7 adjusted to length of 3^rd^ chromosome in a) and the X chromosome in b) and c). Deviations in c) are likely due to uncorrectable viability effects from Sturtevant and Beadle (1936), see Discussion for further explanation.

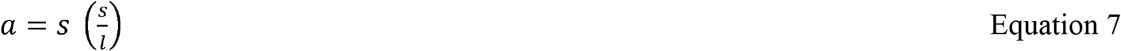

This function is consistent with experimental results of roughly 100% of wildtype crossing over in heterozygotes for the full chromosome inversion *In(1)y^4^* and between 25-50% the wildtype rate for shorter inversions *In(1)sc^7^* and *In(1)dl-49* (Sturtevant and Beadle 1936, Novitski and Braver 1954). The experimental results for reduced crossing over in heterozygous inversions is equivalent to the reduction observed for crossing over in homosequential regions of overlapping inversions (compare figure 2b and 2c). Thus equation 7 can also be used to predict the probability of forming an asymmetric dyads (a) where s is the size of the *shared* inverted region.

Despite the relative obscurity of non-random disjunction in modern genetics, there was an extensive series of experiments estimating the probability of recovery from asymmetric dyads in the golden era of chromosome mechanics (reviewed in Novitski 1967, Lucchesi 1994). This evidence comes from experiments on heterochromatin displacement (Gershenson 1933, Novitski 1951, Novitski 1967), translocation heterozygotes (Glass 1935, Zimmering 1955, Chandley 1965), and numerous estimates from compound chromosomes of all possible configurations (Novitski and Sandler 1956, Sandler 1958, Sandler and Lindsley 1963, Lindsley and Sandler 1965, Lucchesi 1965, Sandler 1965, Sandler and Lindsley 1967). Figure 3 combines all these experiments and reveals a surprisingly basic relationship between degree of dyad asymmetry (*d*) in meiosis II and the probability of recovery from an asymmetric dyad (*r*). This data suggests a simple function

**Figure 3.**
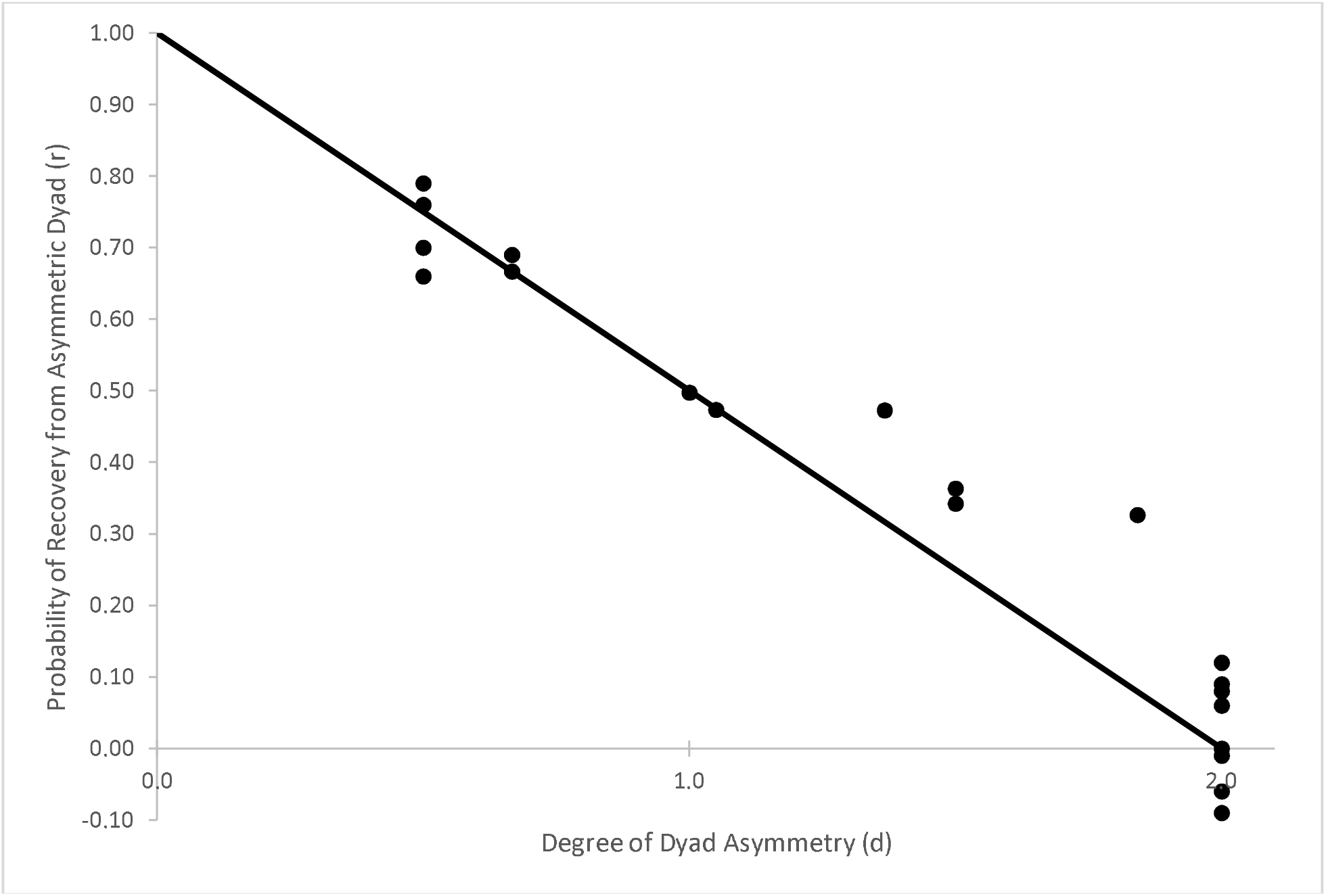
Probability of recovering the shorter chromatid of an asymmetric dyad. A probability less than zero is an artifact of complex viability corrections for whole chromosome duplications rarely included in function egg. Illustrated function is not fitted to data, but rather is the function described in equation 8 for probability of recovering the shorter chromatid of an asymmetric dyad.

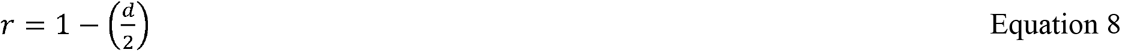

which results in Mendelian segregation for symmetric dyads (*d = 1*) and accurately predicts the complete drive against compound chromosomes twice the size of normal chromosomes (*d = 2*).

As it pertains to crossing over in shared inverted regions of overlapping inversions, the degree of asymmetry is determined by the size of regions duplicated in one recombinant class, which are the same as those deleted in the other recombinant class. Therefore, the asymmetry generated in each dyad is roughly equivalent, especially for small deletions and duplications. However, degree of asymmetry is a relative measure which means a deletion creates a slightly greater asymmetry than the equivalent size duplication, as 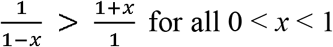. This second form of asymmetry, *among* rather than within dyads, requires separate estimates of *r_prox_* and *r_dist_*. In summary, numerical predictions for nonrandom disjunction in *D. melanogaster* females heterozygous for any two inversions can be generated from the probability of a crossover forming an asymmetric dyad and the probability of recovery from that asymmetric dyad, which in turn are basic functions of the size of the shared inverted region and the degree of the resulting dyad asymmetry, respectively.

### Persistence Time of New Inversions

The survival probability of a new mutant is of fundamental importance in population genetics. However, other statistics such as mean time to extinction (persistence) or the total number of individuals carrying the mutation before extinction (pervasiveness), may be of greater practical importance (Garcia-Dorado, Caballero et al. 2003). In this section the survival probabilities (equations 4 a,b,c,d) are used to calculate persistence for all possible inversions entering the already polymorphic natural populations of *D. melanogaster*. In the following section the pervasiveness for these same inversions are calculated to demonstrate that those gene arrangements actually observed in natural populations of *D. melanogaster* are not a random sample of all possible inversions. These calculations illustrate how small biases in the transmission (Δ*k* ≤ 0.015) and slight alterations of survival probabilities (Δ*u*_1_ ≤ 0.025) for chromosomal rearrangements can produce major patterns in inversion polymorphism.

The *Drosophila melanogaster* karyotype consists of one acrocentric sex chromosome (Muller Element A), two metacentric autosomes (Muller Elements B, C, and D, E, respectively), and a heterochromatin rich dot chromosome (Muller Element F) (Muller 1940). The study of chromosomal rearrangements in this species represents the largest and most thorough catalogue of paracentric inversions (Lemeuenier and Aulard 1992). Each chromosome arm of the metacentric autosomes consists of 120 cytological subdivisions and each arm has a large, subtelomeric paracentric inversion that is found in all natural populations worldwide at frequencies between 0.05 to above 0.50; the so-called common cosmopolitan inversions (Lemeuenier and Aulard 1992). These inversions are young relative to the species split with *D. simulans* (~5 mya), but are older than *D. melanogaster’s* worldwide expansion out of sub-Saharan Africa (~100 kya) (Wesley and Eanes 1994, Andolfatto et al. 1999, Tamura et al. 2004, Matzkin et al. 2005, Corbett-Detig and Hartl 2012). In addition to these four common cosmopolitan inversions [*In(2L)t, In(2R)NS, In(3L)P*, and *In(3R)P*] there are 31 rare, but repeatedly observed inversions at frequencies below 0.05, as well as 283 unique inversions that have been observed only as a single copy in samples from natural populations. In contrast, no inversions have ever been observed in the dot chromosome, and inversions of the acrocentric X chromosome are fewer with no populations outside of Africa having inversions segregating at high frequency (Lemeuenier and Aulard 1992).

Despite the extensive effort focused on *D. melanogaster*, this catalogue represents only about 1% of all possible paracentric inversions in this species. Given the cytological limitations of detection “all possible inversions” are any gene rearrangements spanning at least one cytological subdivision yielding 7140 possible paracentric inversions per chromosome arm. Because these largely unobserved spontaneous rearrangements occur in populations that invariably have a large, subtelomeric inversions segregating at high frequency, the survival dynamics of newly inverted chromosomes are subject to the cytogenetic mechanism described in this paper.

For every possible inversion the probability of forming an asymmetric dyad (a) was calculated using equation 7 with genetic lengths based on the cytological to genetic map conversion for *D. melanogaster* downloaded from Flybase March 1^st^ 2017. The degree of dyad asymmetry (*d*) was calculated based on the number of 120 cytological subdivisions per arm that are either duplicated (*dup*) or deleted (*del*) due to crossing over in the shared inverted regions, which itself varies depending on relative size and position of the new inversion

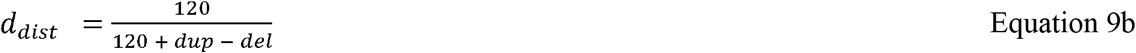

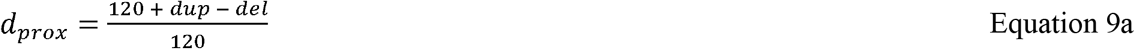

The degree of dyad asymmetry (*d*) was converted to probability of recovery from asymmetric dyad (*r*) using equation 8. These values for crossing over and disjunction were then entered into equation 6 (a,b,c,d) using cosmopolitan inversion frequency (*q*) of 0.25 to generate the survival probability for every possible inversion entering a natural population of *D. melanogaster*. The mean persistence time 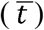 for each newly inverted chromosome is the sum of the infinite series based on the single generation survival probabilities

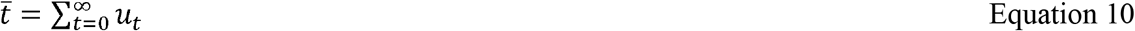

The limit for the series was estimated based on partial sums by numerical iteration for 10,000 generations.

Twenty percent of all possible paracentric inversions will not overlap the large, subtelomeric inversions found on each chromosome arm. Independent inversions, *i.e*. gene rearrangements that do not have any shared inverted regions with the common cosmopolitan inversions, are not subject to the cytogenetic mechanism described in this article and in an infinite population have an unlimited persistence time. In finite populations, the cumulative number of copies for neutral variants is 2*N_e_* (Maruyama 1971), which far exceeds to the numerical values for any overlapping inversions (see next section for the expected number of copies calculation). Considering inversions observed at least once in natural populations, rather than independent inversions being more numerous than overlapping inversions, they were actually underrepresented, making up only 17% instead of the expected 20%. If this analysis is limited to only repeatedly observed gene rearrangements, independent inversion only make up 16%, indicating the bias favoring inversion overlap is even stronger. Clearly, gene rearrangements that share inverted regions with common cosmopolitan inversions are being sampled more often than expected in natural populations. This sampling bias could be caused by the meiotic drive mechanism in absence of strong underdominance or an unobserved mutational biases.

Among overlapping inversions, *i.e*. gene rearrangements that have shared inverted regions with the common cosmopolitan inversions, the cytogenetic mechanism has a disproportionate effect on larger, proximal inversions. To illustrate this bias, the persistence times for all possible inversions were plotted as a function of inversion size and position (figure 4). Inversion size and position was measured on a cytological scale ranging from 0 at the centromere to 120 at the telomere, with each unit representing a subdivision of a cytological band. On this scale inversion size is the distance between breakpoints and position is the value at cytological midpoint of the inversion. Plotting persistence times as a function of inversion size and position produces an uneven “survival surface” for inversions arising on each standard chromosome arm resulting from presence of the large, subtelomeric inversion segregating at high frequency in that population (Figure 4 a,b,c,d,e). As stated earlier, the strength of transmission biases is altered based on the linkage phase of gene rearrangements, therefore the persistence times were calculated for new inversions arising in coupling phase with the common cosmopolitan inversions.

**Figure 4.**
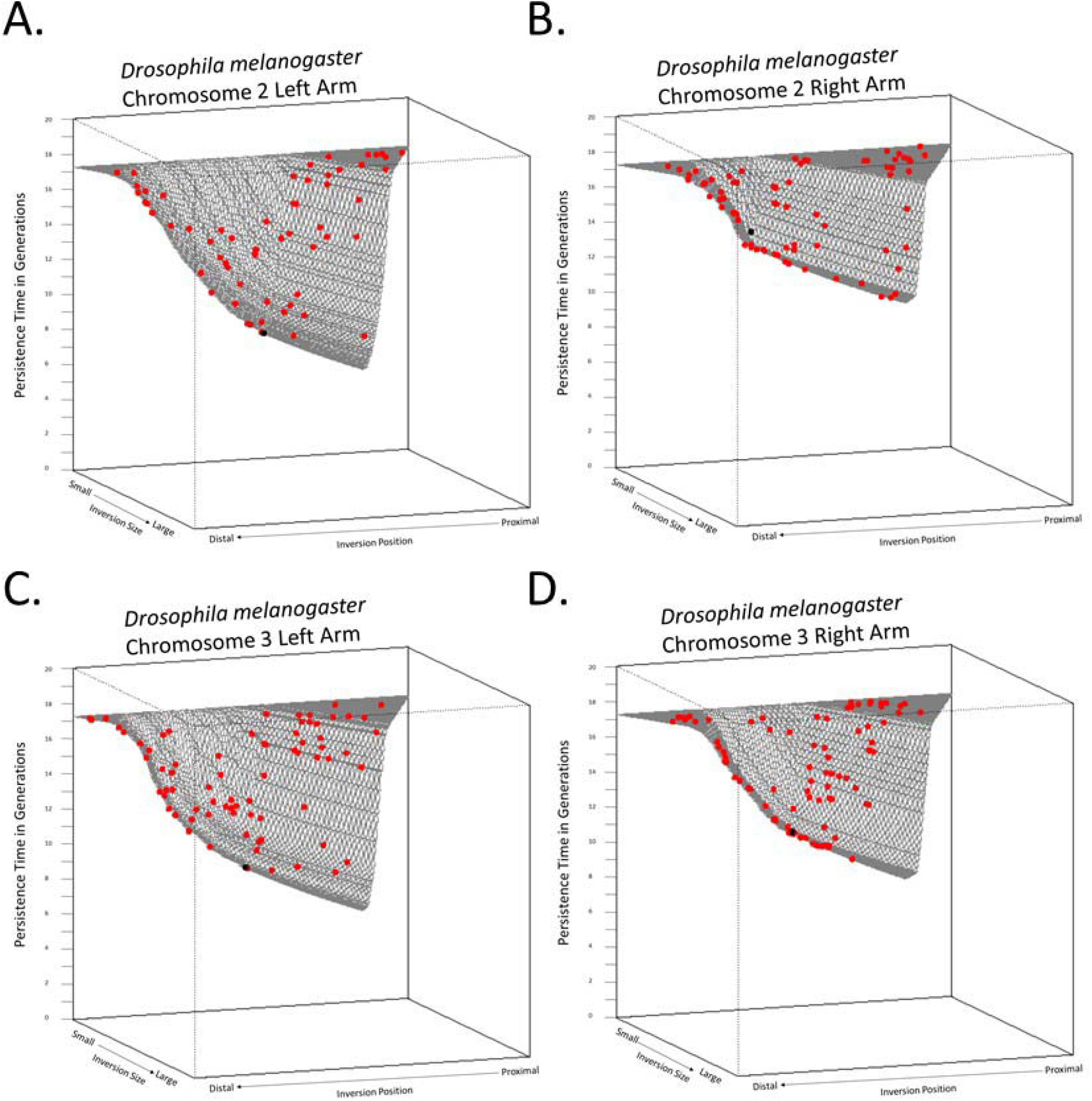
Persistence times for all possible paracentric inversions for a) chromosome two left arm, b) chromosome two right arm, c) chromosome three left arm, and d) chromosome three right arm. The size and position of the common cosmopolitan inversions *In(2L)t, In(2R)NS, In(3L)P*, and *In(3R)P* respectively are illustrated in black. The size and position of all observed inversions from natural populations are illustrated in red.

These plots illustrate all new inversions in natural population of *D. melanogaster* do not have similar persistence times, and those few rare inversions we observe are not randomly distributed with respect to size, position, or linkage phase (Figure 4). Visual inspection of the survival surfaces reveal several “gaps” in the joint distribution of observed inversion size and position. The most noticeable and consistent deficit is seen for the rarely observed large, proximal, repulsion phase inversions and the general absence of coupling phase inversions in *D. melanogaster*. Rather than explain this absence as an unobserved mutational bias or the result of unspecified natural selection, the cytogenetic mechanism presented in this paper predicts their rarity based solely on the probability of forming an asymmetric dyad via crossing over in meiosis I and the probability of recovery from an asymmetric dyad during disjunction in meiosis II.

### Pervasiveness in Natural Population

Expectations for mean time to extinction (persistence) concisely summarizes the population genetic dynamics for new inversions entering natural populations of *D. melanogaster*. Few populations, if any, are sufficiently monitored to capture this dynamic. Fortunately, the survival probabilities can also be used to calculate the cumulative number individuals carrying a newly inverted chromosome before it goes extinct (pervasiveness), giving a relative measure of the probability of detecting inversions of particular size, position, and phase. By substituting the pertinent terms of the cytogenetic mechanism (*ck* = 1 − *hs*) into Garcia-Dorado et al. (2003) branching process derivation for pervasiveness 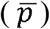 in an infinite population, the expected cumulative number of copies of an newly inverted chromosome before being lost from the population is

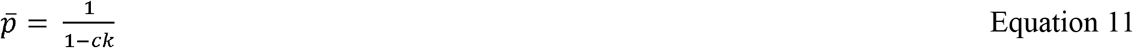

Following the method of the previous section, 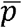 was calculated for all possible inversions. The distribution of pervasiveness for all possible inversions was compared to the distribution of expected pervasiveness for every recorded inversion found in samples from natural populations of *D. melanogaster* (Figure 5). These inversions are subdivided into those observed only once (unique endemics) and those inversions repeatedly observed (recurrent endemics and rare cosmopolitans) (Lemeuenier and Aulard 1992).

**Figure 5.**
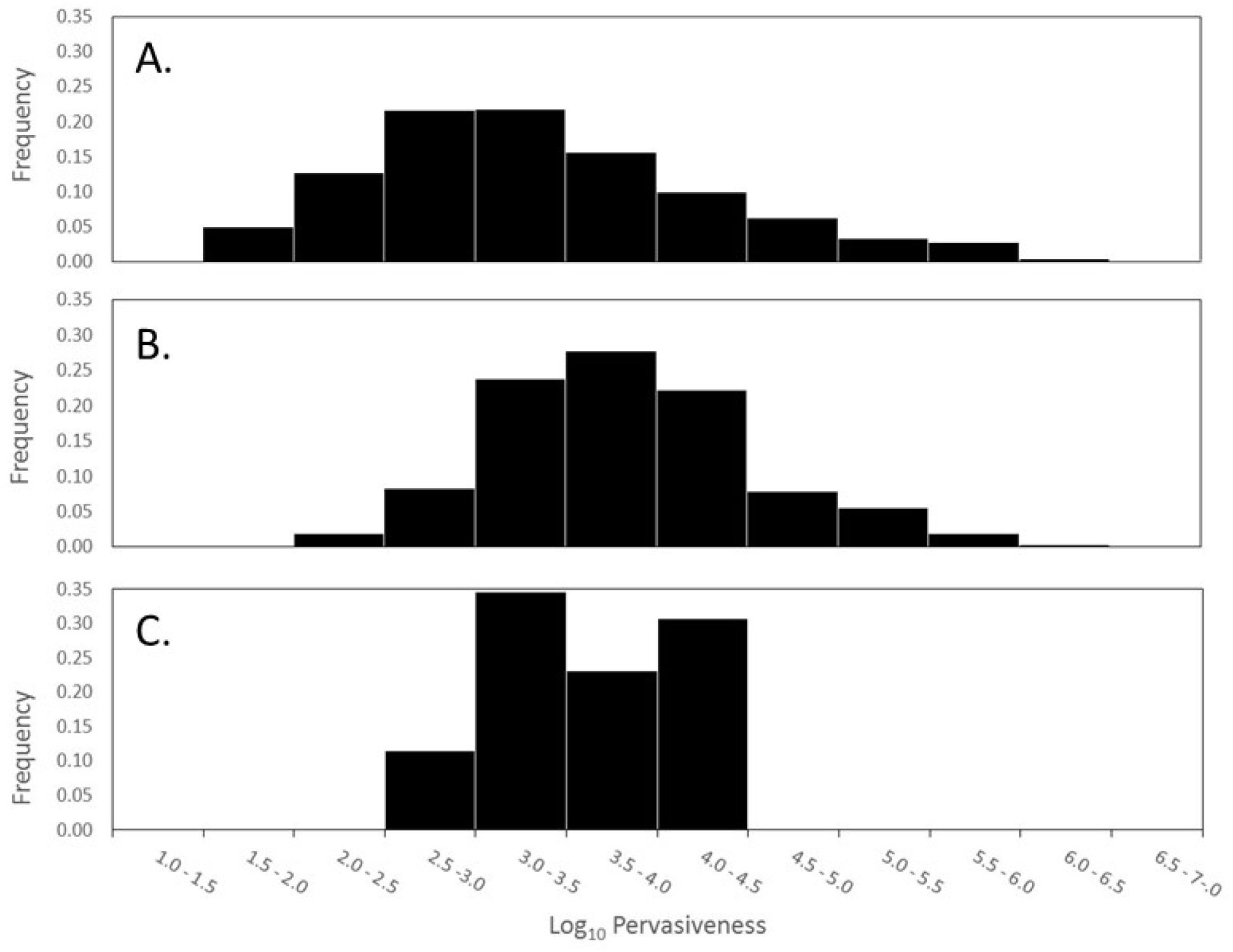
Frequency distribution of expected pervasiveness for a) all possible inversions, b) inversions observed once, and c) inversions repeatedly sampled from natural populations. Because there is over six orders of magnitude difference in expected pervasiveness, the bins of histogram are made on the log10 scale.

Because pervasiveness is a much more sensitive metric than persistence time, this cytogenetic mechanism alone creates a six order of magnitude difference in the expected number of copies before extinction. The 1% of inversion observed at least once is not a random sample of all possible inversion based on pervasiveness (Kolgomorov-Smirnov *D* = 0.318, *p* < 0.001) nor is the 0.1% of inversions that have been recurrently sampled (Kolgomorov-Smirnov *D* = 0.357, *p* = 0.003). This difference is due to the observed inversions having higher expected pervasiveness, while those inversions with predicted to have a lower pervasiveness 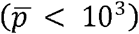 in the presence of the cytogenetic mechanism are largely absent in the catalogue of all known inversions from natural populations (Figure 5b).

## Discussion

New gene arrangements are at constant risk of stochastic loss from populations, irrespective of their fitness effects (Fisher 1923, Haldane 1927, Crow and Kimura 1970). For new inversions entering a population already polymorphic for chromosomal inversions the risk of stochastic loss is dependent on the relative size, position, and linkage phase of the gene rearrangements. Using the branching process, this risk was quantified with a recursion equation for survival probability that incorporates transmission bias and underdominance for all possible inversions entering a natural population of *D. melanogaster*. Persistence times and pervasiveness based on this survival probability demonstrates a distally positioned inversion is expected to have up to 1000-fold more copies over its lifetime than a proximally placed inversion of the same size and fitness. Comparing equivalent inversions also revealed biases favoring smaller inversions (up to 70-fold) and inversions in repulsion phase (7-fold). As expected, inversions observed in natural populations of *D. melanogaster* tend to be smaller, more distal, and in repulsion phase with common cosmopolitan inversions. Thus, the inversions observed in natural populations of *D. melanogaster* are those expected to have the longest persistence and highest pervasiveness under the cytological mechanism described in this article.

Despite the consistency of this cytogenetic mechanism with previously unexplained patterns in inversion polymorphism, it is not a complete model for inversion evolution. This model is predicated on the existence of common cosmopolitan inversions [*In(2L)t, In(2R)NS, In(3L)P*, and *In(3R)P*], which are invariably observed for all natural populations of *D. melanogaster*. Because the cytogenetic mechanism generates underdominance for *all* inversions in polymorphic populations it should result in the selective elimination of those large, subtelomeric cosmopolitan inversions at high frequency. Expectations for underdominance are not always experimentally fulfilled, as has been thoroughly demonstrated for pericentric inversions in *D. melanogaster*. In this case pericentric inversions that did not exhibit expected underdominance were those whose breakpoints disrupted “sensitive sites” necessary for normal crossing over (Coyne et al. 1993). Interestingly, Corbett-Detig (2016) showed that breakpoints of cosmopolitan and recurrent paracentric inversions in *D. melanogaster* are closer to cytological bands containing sensitive sites than the breakpoints of rare inversions. This pattern was interpreted as evidence of positive selection for recombination suppression *per se*, but upon considering the crossover dependent fertility underdominance generated by overlapping paracentric inversions it would appear common cosmopolitan inversions’ proximity to sensitive sites prevents them from suffering the effects of underdominance when heterozygous with another inversion. Thus the pattern of sensitive site association can be interpreted as the result of stronger purifying selection on fertility effects for rare inversions that accelerates their removal from the population.

If cosmopolitan inversions escape the effect of underdominance by disrupting normal function of sensitive sites, they represent a small exception (0.01 %) to the general rule for chromosomal inversions generally. The underdominance associated with chromosomal rearrangements is well-supported by the relative rarity of segregating translocations and pericentric inversions despite their occurrence as fixed differences between species (White 1977, Hooper and Price 2015). Furthermore, underdominance for overlapping paracentric inversion has been explicitly demonstrated by Sturtevant and Beadle (1936) where only 5 of 16 overlapping inversion pairs had any viable crossover progeny, and in each of those 5 combinations at least one of the meiotic products was dominant sterile or lethal. The fact that both duplications and deletions exhibit partial viability effects, which are usually more severe for deletions, is likely the cause of low estimates of crossing over in shared inverted regions observed in figure 2c. Despite the theoretical, comparative, and experimental consensus on underdominance of gene rearrangements, there is an equally large body of evidence from natural populations and laboratory demonstrations supporting the selective maintenance of paracentric inversions (e.g. Wright and Dobzhansky 1946, Schaeffer 2008). This contradiction presents a major problem for any general or complete model of inversion evolution.

There are three phases in the life of any mutant; origin, establishment, and ultimate loss or fixation. The mechanisms and balance of forces governing allele frequency change in each phase are not necessarily the same. Generally, stochastic processes gain importance at extremely low or high frequency, whereas selection predominates at moderate frequency (Crow and Kimura 1970). Chromosomal rearrangements are a special case because balancing selection appears to maintain inversions at appreciable frequencies and delays the approach to their ultimate fate. Much of theory has focused on selective mechanisms (coadaptation and local adaptation) to explain the behavior of these common inversions (Kirkpatrick and Barton 2006, Hoffmann and Rieseberg 2008). However, in *D. melanogaster* these large, subtelomeric inversions maintained by balancing selection represent approximately 1% (4/318) of observed inversions, and only 0.01% (4/28560) of all possible paracentric inversions. The present study focuses on predominantly non-adaptive mechanisms for the short-lived and rarely (or never) observed inversions. In fact, the pervasiveness calculations give a precise quantification of the ascertainment bias involved in cataloguing the rare endemic inversions of *D. melanogaster*.

Although processes described here have proven useful in explaining patterns in observed inversion polymorphism of *D. melanogaster*, it is important to note that both rare and common gene rearrangements show biases toward smaller, distal inversions. Similarly, in *D. pseudoobscura* this cytogenetic mechanism is thought to produce a distal shift and size reduction of inversions in a phylogenetic series, but rare and common gene arrangements contribute equally to this pattern. This equal contribution is difficult to explain because the proposed meiotic drive mechanism cannot alone cause inversions to increase to high frequency, and there is no necessary reason that common, selectively maintained inversions should conform to the patterns observed for rare inversions.

This apparent paradox can be solved by shifting focus to the meiotic drag of larger, proximal inversions. These inversions are rapidly eliminated from populations because they suffer from the compounded effects of undertransmission and underdominance. Selective advantages as great as 5% are insufficient to overcome this double penalty. Thus, even if proximally located inversions capture locally adapted alleles or beneficial epistatic combinations they will likely be lost, whereas distally located inversions with similar fitness effects persist long enough for selection to take hold and drive their establishment in the population. In this manner the patterns generated by non-adaptive mechanisms for rare inversions entering the populations are translated to patterns of common inversions maintained at appreciable frequencies by balancing selection. It should come as no surprise then that all four common cosmopolitan inversions in *D. melanogaster* are subtelomeric (distally positioned) and the more recently favored gene arrangements in inversion laden species of the *obscura* group species all show the distal shift.

## Conclusion

The complex meiotic behavior of overlapping inversions is predicted to alter the survival probability of newly inverted chromosomes. This is because females heterozygous for overlapping inversions will exhibit fertility underdominance and meiotic drive/drag as a consequence of asymmetric dyad formation in meiosis. Survival probabilities can, therefore, be expressed in terms of probability of forming asymmetric dyads and the probability of recovery from those asymmetric dyads. Survival probabilities and the related population genetic statistics of persistence and pervasiveness, were calculated for all possible inversions entering a natural population of *D. melanogaster* revealing a bias towards the rapid elimination of larger, proximal inversions especially those in coupling phase with cosmopolitan inversions. Consistent with these predictions, the historical record of observed inversions in *D. melanogaster* natural populations were shown to be a non-random sample from all possible inversions and biased towards detecting inversions (smaller, distal, repulsion phase) that are expected to possess the greatest number of copies before going extinct.

## Supporting information

## List of Variables

*m*: = number of copies of a new mutant in the first generation after mutation
*c*: = average family size
*n*: = family size
*k*: = transmission ratio
*t*: = generations after spontaneous mutation
*u_t_*: = probability of survival to generation t
*a*: = probability of forming an asymmetric dyad via crossing over
*r*: = probability of recovering shorter chromatid from an asymmetric dyad
*s*: = size of shared inverted region
*l*: = total genetic length of a chromosome
*d*: = degree of asymmetry in an asymmetric dyad
*q*: = frequency of common cosmopolitan inversion
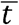: = persistence, average number of generation after spontaneous mutation before extinction
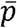: = pervasiveness, cumulative number of copies after spontaneous mutation before extinction

