## Supplementary material for "Meiotic Drive and Survival Probability of Newly Inverted Chromosomes"

Supplemental Figure 1a. Pairing diagrams for overlapping inversions in coupling phase.

Meiotic products for crossover located in Region B, size asymmetry illustrated is proportional.

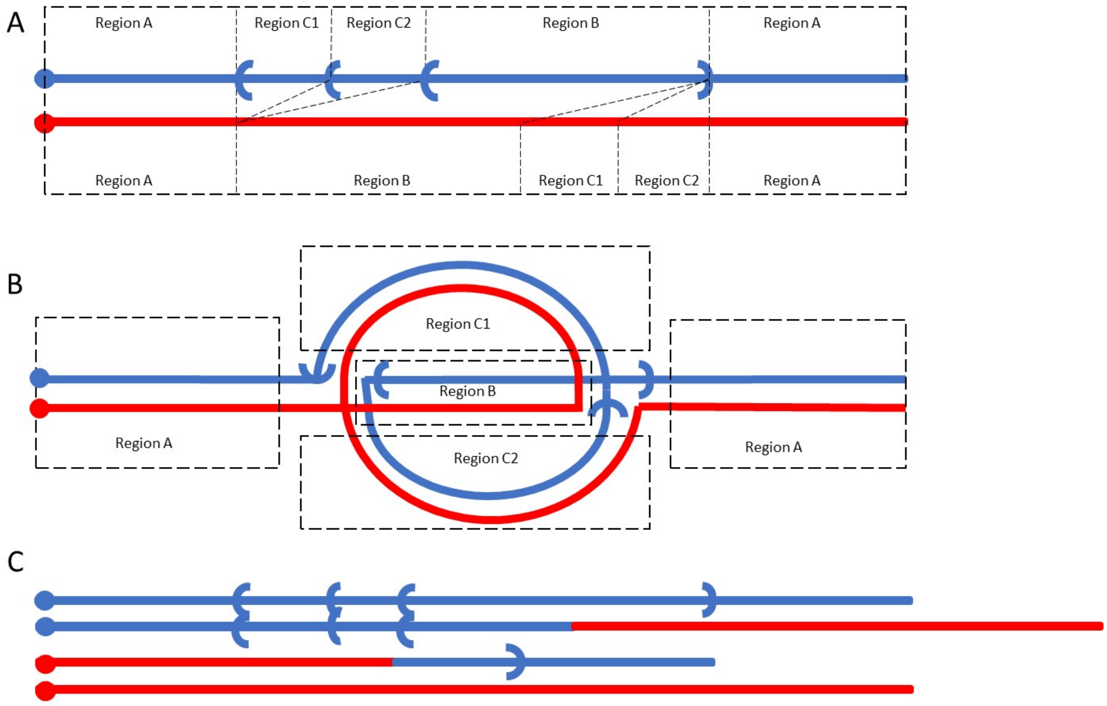

Supplemental Figure 1b. Pairing diagrams for included inversions in repulsion phase.

Meiotic products for crossover located in Region B, size asymmetry illustrated is proportional.

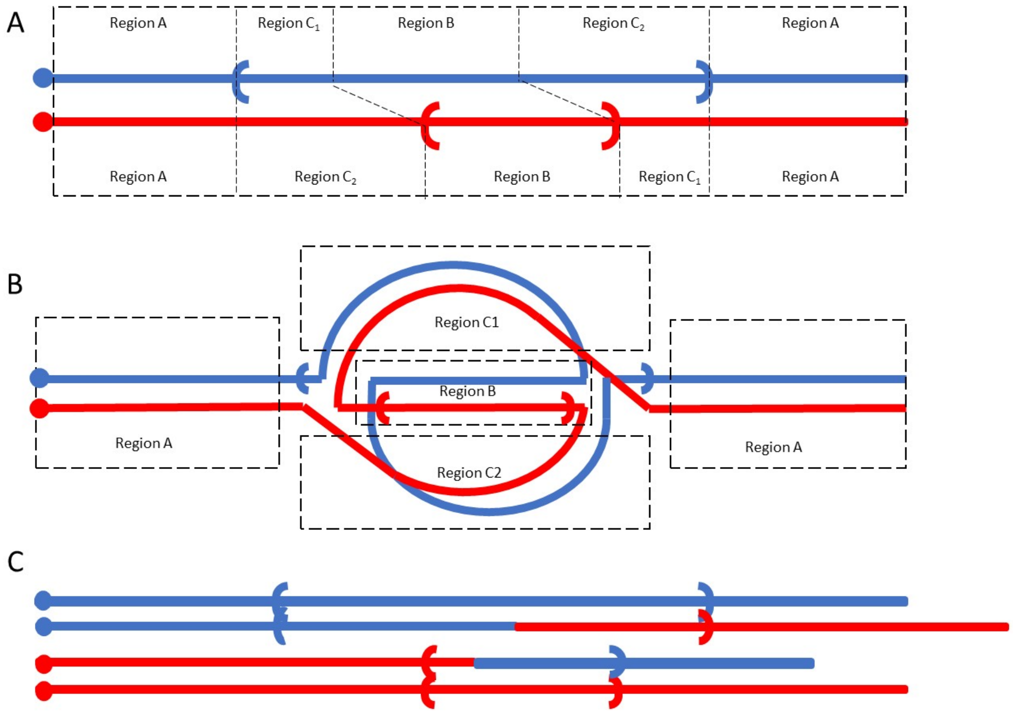

Supplemental Figure 1c. Pairing diagrams for included inversions in coupling phase.

Meiotic products for crossover located in Region B, size asymmetry illustrated is proportional.

A

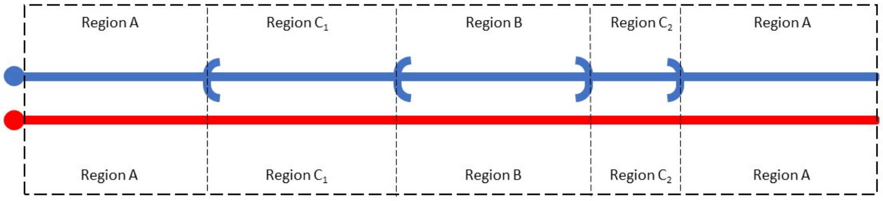


B

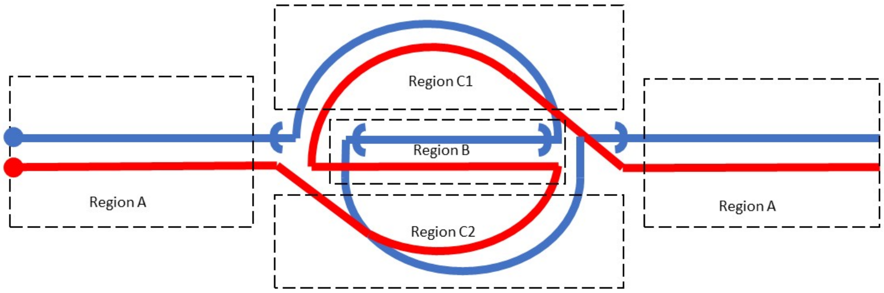


C

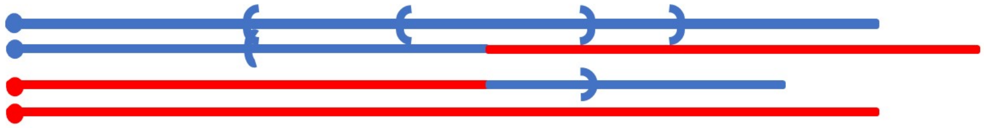
